# Belgian endive-derived biostimulant activity in Arabidopsis, lettuce and sweet pepper at different developmental stages, environmental conditions, and application methods

**DOI:** 10.64898/2026.04.15.717650

**Authors:** Halimat Yewande Ogunsanya, Christophe Petit, Kris Audenaert, Noémie De Zutter, Danny Geelen

## Abstract

Belgian endive-derived biostimulant (BEE) was previously shown to enhance root and shoot growth of *Arabidopsis thaliana* and *Plectranthus esculentus* in *in vitro* culturing conditions. In this study, we evaluated the effect of BEE on *A. thaliana* subjected to abiotic stresses and assessed the translatability of its bioactivity on lettuce (*Lactuca sativa*) and sweet pepper (*Capsicum annuum*) cultured in substrate and soil. A first set of experiments tested the impact of BEE on protection during, and restoration after, osmotic or salt (NaCl) stress. BEE treatment had little to no rescuing effect when plants were exposed to osmotic stress. In contrast, BEE strongly promoted shoot development and leaf health both under standard and NaCl stress conditions. Under mild stress, BEE enhanced photosynthetic efficiency and chlorophyll content in *Arabidopsis*, whereas it did not significantly alleviate osmotic stress induced by sorbitol. To evaluate the effect under *ex vitro* conditions, BEE was applied via root drenching to substrate-grown *A. thaliana,* lettuce, and sweet pepper. BEE improved leaf greenness and photosynthesis enhancing *Arabidopsis* rosette development, but it did not increase lettuce head weight. In sweet pepper, BEE increased fruit yield and promoted fruit maturation. Under drought stress conditions, BEE application did not improve sweet pepper yield.

## 1.0 Introduction

The increasing global population, projected to reach approximately 9.7 billion by 2050, is placing pressure on agricultural systems to meet the growing demand for food. This pressure necessitates innovative and sustainable approaches to enhance agricultural productivity while minimizing environmental impact. One of the promising strategies to achieve this goal is the discovery, development, and application of plant biostimulants.

The discovery of biostimulant activity through screening of (organic) materials often involves treatment of *in vitro* cultured Arabidopsis seedlings followed by validation bioassays (Li et al., 2026). Such studies, however, do not provide insight into putative biostimulant activity in horticultural plants, nor whether activity is preserved under field or substrate cultivation conditions. To evaluate the potential of a biostimulant, studies using economically important species (e.g. sweet pepper) cultivated under conditions are necessary (Calvo et al., 2014). Additionally, the effectiveness of biostimulants is often assessed under different environmental stress conditions, including drought stress, salinity stress, and temperature variations, to determine their adaptability and mode of action in real-world farming systems. Such trials provide valuable insights into dosage optimization, application timing, and potential interactions with conventional agricultural practices, ensuring that the biostimulant is both effective and practical for large-scale implementation.

Salt and drought stress are increasingly important challenges in sweet pepper cultivation under greenhouse conditions, where optimal control of environmental factors is essential for high productivity and fruit quality. Despite the controlled environment, water use efficiency remains a concern, especially as greenhouse agriculture expands in water-limited regions. Drought stress in greenhouses can impair stomatal function, reduce photosynthetic efficiency, and limit biomass accumulation in sweet pepper plants (Dorji et al., 2005). Similarly, salt accumulation in recirculating hydroponic or soilless systems; common in greenhouses, can lead to ionic imbalance, osmotic stress, and nutrient antagonism, ultimately affecting fruit development and yield (Savvas & Gruda, 2018). Plant extract biostimulants have been reported to be very effective in promoting growth in field experiments (Li, Van Gerrewey, et al., 2022). The impact of biostimulant application is often most pronounced under conditions of stress (Calvo et al., 2014; Colla & Rouphael, 2020; du Jardin, 2015).

Biostimulants may enhance vegetative and or reproductive stages (Parađiković et al., 2019). While many commercially available biostimulants exhibit activity across multiple growth stages, identifying the specific developmental stage targeted by a biostimulant can enhance its efficiency. For instance, biostimulants that specifically promote vegetative growth can be employed during early crop establishment to ensure strong root and shoot development, while those that enhance reproductive growth can be applied during flowering and fruiting stages to improve yield and quality. Likewise, biostimulants that enhance vegetative growth may particularly promote growth of leafy vegetables like lettuce, while those that promote reproductive development may be more suitable for fruit-bearing crops like tomatoes and sweet pepper. This stage-specific application of biostimulants can lead to more efficient agricultural practices by maximizing crop performance and yield.

Previously, we reported on *in vitro* root and shoot growth stimulating activity of Belgian endive extracts (BEE) derived from the forced roots that are a waste product from Belgian endive production (Ogunsanya et al., 2022). In our follow up study, we analysed the impact of BEE across different environmental conditions using the model plant *A. thaliana* and agricultural-relevant crop plants, lettuce and sweet pepper. These experiments included *in vitro* assays assessing the physiological effects, as well as indoor and greenhouse experiments. The efficacy of BEE under standard growth conditions and abiotic stress conditions, including salinity stress and drought stress, both of which represent critical challenges to global agricultural productivity was assessed. Additionally, while a previous study has highlighted the positive effect of BEE on vegetative growth *in vitro* (Ogunsanya et al., 2022), we here explore the impact of BEE on both vegetative and reproductive growth *ex vitro*. Our findings reveal a highly complex mode of action of BEE on both plant growth and development.

## 2.0 Materials and Methods

### 2.1 Seed sterilization, media composition and growth conditions

*Arabidopsis thaliana* Col-0 wildtype seeds were vapor-phase sterilized and sown on 12 x 12 cm^2^ plate containing Murishige and Skoog (MS) medium. The MS medium consisted of 1.5 g/L MS basal salt, 0.5 g/L MES buffer (2-(N-morpholino)ethanesulfonic acid), 5 g/L sucrose, 8 g/L plant agar, and adjusted to pH 5.7 with 1M KOH. Seeds were subjected to stratification in the dark at 5°C for four days to ensure synchronized germination, followed by an 8 h light exposure in the growth room (100 µmol/m^2^/s intensity, 16 h light/8 h dark photoperiod at 21 °C) and subsequently kept in the dark for 3 days to induce etiolation (Trinh et al., 2018). Seeds transferred to growing substrates did not undergo etiolation and was kept in light for 5 days.

### 2.2 Osmotic stress via root assay

After three days of etiolation, Arabidopsis seedlings were transferred to treatment plates. The treatment plates included either mock MS medium (no sorbitol), MS supplemented with 100 mM sorbitol, or MS supplemented with 150 mM sorbitol. Each stress condition was combined with either 0.36 g/L BEE, 0.71 g/L BEE, or no BEE, as detailed in Table 2.1. Ten days after transfer, seedlings were imaged for phenotypic analysis. Primary root length and leaf area were measured using Fiji (ImageJ), while lateral root numbers were counted under a binocular microscope (Olympus SZX9).

**Table 2.1.** BEE-treatments included in the osmotic stress-assay on *Arabidopsis thaliana* Col-0 seedlings, induced by different sorbitol concentrations. BEE, Belgian Endive Extract.

| Treatments | BEE (0 g/L) | BEE (0.36 g/L) | BEE (0.71 g/L) |
| --- | --- | --- | --- |
| **Sorbitol (0 mM)** | Standard Control | Extract Control 1 | Extract Control 2 |
| **Sorbitol (100 mM)** | Positive Control 1 | Test 1 | Test 2 |
| **Sorbitol (150 mM)** | Positive Control 2 | Test 3 | Test 4 |

Sorbitol (0 mM) = growth medium only without sorbitol. BEE (0%) = growth medium only without BEE.

### 2.3 Salinity stress via true leaf development assay

The true leaf development assay was adapted from (Li, Evon, et al., 2022) to evaluate the effects of BEE on early vegetative growth under salt stress. *Arabidopsis thaliana* seeds were pre-selected using binocular microscope (Olympus) to ensure viability and reduce variability. Sterilized seeds were then sown on MS medium supplemented with either 0 mM or 100 mM NaCl, with or without 0.1 g/L of BEE. The 100 mM NaCl concentration was used to induce mild salinity stress. Following sowing, seeds were stratified at 5°C in darkness for four days, after which they were incubated horizontally in a growth room (100 µmol/m^2^/s light intensity, 16 h light/8 h dark photoperiod at 21 °C) for 10 days. Each treatment consisted of three replicate plates, with 50 seeds per plate, to further minimize seed-to-seed variability. Germination was assessed by calculating the proportion of seeds showing visible radicle emergence post-testa rupture, relative to the total number of sown seeds. First "true" leaves development was monitored through imaging six days after sowing (DAS) using a binocular microscope (Olympus SZX9) with a 10x eyepiece (WHS10X-H/22) and 1x objective lens (DFPLAPO1x-2). It was used at a zoom setting of 2.0x corresponding to a total magnification of 20X. The percentage of true leaf emergence was calculated as the number of germinated seeds that developed true leaves, divided by the total number of germinated seeds. Ten days after sowing, plants images were captured with a RGB Nikon camera, and leaf area was quantified using ImageJ. To complement morphological observations, plant health and development were evaluated using a multispectral imaging platform (Phenovation, Wageningen, The Netherlands). The overall plant health status was evaluated using the chlorophyll fluorescence (Fv/Fm), which is a proxy for the efficiency of photosynthesis (Baker, 2008). Chromophore-based proxies were included to evaluate the chlorophyll and anthocyanin content of the Arabidopsis plants, and more specifically the chlorophyll index (ChlIdx,(Gitelson et al., 2003)) and modified anthocyanin reflectance index (mARI, (Gitelson et al., 2009)) were calculated. Detailed descriptions of these spectral parameters are provided in Table S1 of the supplementary materials.

### 2.4 Arabidopsis growth on substrate, treatments, and measurements

*Arabidopsis thaliana* Col-0 wildtype seeds were germinated as described in section 2.1. Next, seedlings were transferred to the Jiffy peat pods (Jiffy Products International AS, Norway) and covered with plastic film for 3 days. The seedlings were treated and left to grow in the growth room (16/8 hours light/dark light, light density 70 µm m^-2^ s^-1^, 21 °C). Two forms of BEE treatment were used: heat-treated (autoclaved, Auto_BEE) or not (non-autoclaved, NA_BEE). The treatments were further applied in two ways; root drenching and shoot spraying. Each treatment contained 18 plants. Respective treatments i.e., (non)autoclaved BEE at 0.18 g/L, 0.36 g/L, and control were applied to the plants 5 days after transfer to the Jiffy pods. The root treatment was achieved by drenching the tray containing the plants in the respective treatment solutions (∼1-2 L). The shoot treatment was achieved by spraying each plant with ∼3 mL of respective treatments 5 days after transfer to the jiffy pods (i.e., 14 days after sowing). The plants were subsequently watered twice to thrice a week with fertilizer (Wuxal 8-8-6).

The shoot area was measured as a vegetative parameter. Fourteen days after germination (DAG), the shoot phenotype of each plant was evaluated. A Nikon RGB- camera was used to take pictures of the trays containing the plants for leaf area scoring. The scoring of the leaf area was done using Fiji.

### 2.5 Lettuce growth, treatments, and measurements

Lettuce (*Lactuca sativa* var. Expertise, Rijk Zwaan) seeds were germinated directly on pre-wet Jiffy peat pods and kept in a growth chamber under controlled conditions: 18 °C, 60-70% relative humidity, and 200-220 μmol/m⁻ ²/s light intensity with a 16-hour photoperiod. Uniformly grown seedlings were selected and transferred into new trays 10 days after sowing. Six seedlings were selected per tray per treatment with 3 technical replicates. The lettuce seedlings were treated with 0.18 g/L, 0.36 g/L of BEE or not (Control). Plants were evaluated 41 days after sowing. Fresh weight of the lettuce shoot was quantified after which they were dried in a n oven at 70 °C for 24-48 hours for dry weight quantification. PSI Polypen was used to measure the leaf indices. The Normalized Difference Vegetation Index (NDVI), Greenness Model 1 (GM1), Carotenoid Reflectance Index 1 (CRI1), Anthocyanin Reflectance Index 1 (ARI1), General coefficient (G), Control Value 1 (Ctr1) were recorded.

### 2.6 Sweet pepper growth, treatments, and measurements

The sweet pepper cultivar *Capsicum annuum* ‘MADURO F1’ (F1 hybrid) seeds used were sourced from Enza Zaden (Enkhuizen, The Netherlands). Sweet pepper seeds were put in 40 °C water to break dormancy, after which they were germinated on wet Whatman paper in a petri dish. The petri dish containing the seeds was wrapped in aluminium foil and placed in 25 °C growth room in the dark for 3 days. Germinated seedlings were transferred to Jiffy peat pods (Jiffy Products International AS, Norway) and incubated in a growth room under 100-110 µmol/m^2^/s light and 16h/8h photoperiod at 25 °C for 25 days. They were subsequently transplanted into pots containing potting soil or rockwool blocks (depending on the experiment, Table 2.3) and transported to a greenhouse or growth room under controlled temperature, light, and humidity (25-28 °C, 220-260 µmol/m^2^/s, ∼60% respectively). The plants were uniformly irrigated and kept under a 16 h photoperiod. The experiment was conducted with two main groups: BEE group and control group. In the BEE group, the sweet pepper plant received BEE treatment as supplement, while the control group was not supplemented but received water instead. A concentration of 2 g/L of BEE was applied to the plants and a volume 50 mL per plant was given (Table 2.2). The BEE treatment was applied via root application and was done one week before projected flowering. The control plants were given water without supplements. In drought stressed plants, the automatic irrigation of the plants was stopped for 3-5 days until the plants were slightly wilted and the water content in the soil reached about 30 %. Treatment was applied once the induced drought was stopped.

**Table 2.2.**
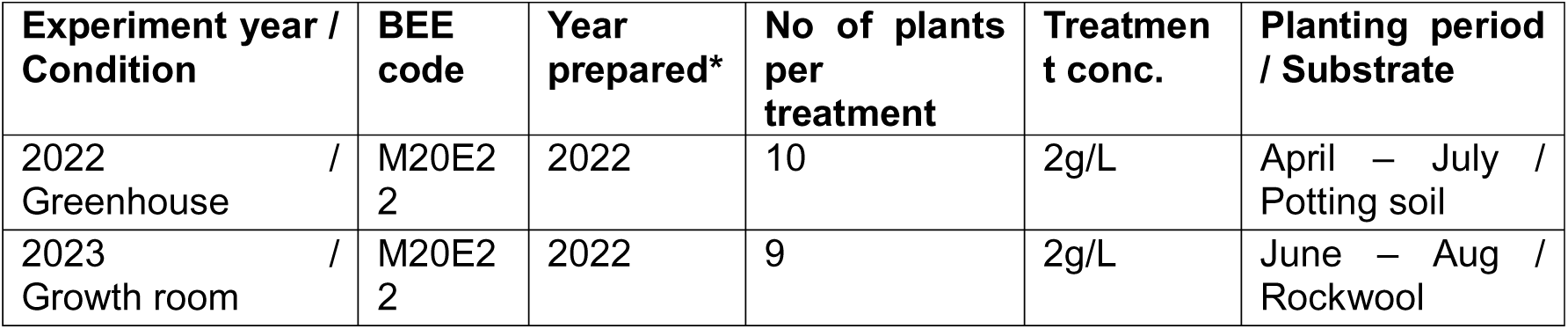
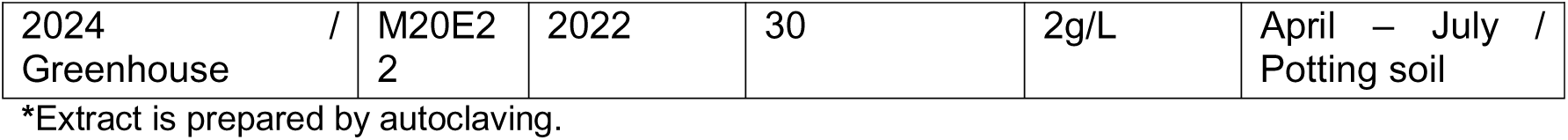
Experimental setup of sweet pepper experiment, detailing the experimental year, BEE treatment, planting season and substrate.

Data were collected to evaluate the effect of BEE on sweet pepper plants on important growth parameters and yield components. These include plant height (cm), chlorophyll content, number of fruits per plant, and fruit yield. The plant height was measured from the base to the top of the plant. The chlorophyll content was measured using a DUALEX meter (ForceA, France) to estimate leaf greenness and photosynthetic potential. The DUALEX measures the light transmitted through the leaf at specific wavelengths to estimate the chlorophyll content as an absolute value in µg/cm^2^. Three young leaves from a branch of each plant were measured and each leaf was read at 3 different spots. The number of fruits per plant was counted to evaluate the effect of BEE on fruit production. The total fruit yield per plant was recorded as the fresh weight of all fruits (in g) harvested at the end of the experiment. All data were collected at the end of the experiment.

### 2.7 Ethylene measurements

The release of ethylene was measured in sweet pepper plants 12 hours, 24 hours, and 48 hours after treatment with BEE. A control measurement (0 hour) was measured before the BEE treatment. A young leaf per plant of sweet pepper was detached and rolled into a 7 mL glass vial (Supelco). The glass vial was left open for 15 minutes to evaporate the wound-induced ethylene burst released from detaching the leaves. Next, the glass vials were closed with screw caps equipped with PTFE/silicone septum (Supelco) and incubated for 4 hours at 25 °C in the dark for ethylene accumulation in the headspace of the vial. After incubation, 2 mL of the headspace was taken using a gastight syringe (Supelco) and analyzed using a gas chromatography (GC, Trace 1300, Thermo Scientific Waltham, MA, USA) equipped with a flame ionization and detector. Calibration was done using standards of 0.4 PPM (parts per million) and 1 PPM ethylene. Subsequently, the leaves in the vial were dried in the oven (80°C) and their weight was used in calculating the ethylene content.

### 2.8 Statistical analyses

Data manipulation and visualization were performed using Python (version 3.2). The pandas library (Mckinney, 2011) was used for data preprocessing, while NumPy (Harris et al., 2020) was used for numerical computations. Data visualization was carried out using Matplotlib (Hunter, 2007) and Seaborn (Waskom, 2021). Data were visualized using boxplots with individual datapoint overlaid. Boxes represent the interquartile range (IQR) with the median, and whiskers indicate the most extreme values within 1.5 x IQR from Q1 and Q3. For datasets with a low number of observations, bar plots displaying the mean ± standard deviation were used instead. Bar plots were also utilized for all graphs within figures where the use of boxplots was not appropriate. Normality and homoscedasticity test were conducted using Shapiro-Wilk and Levene’s tests respectively to check if the data were normally distributed. Data that failed the normality test were subjected to non-parametric testing using Krustal-Wallis test, otherwise, a parametric testing using ANOVA (Analysis of Variance) was used. Statistical analyses, including ANOVA and t-test, were conducted using SciPy (Virtanen et al., 2020) and/or Pingouin (Vallat, 2018). A value of p < 0.05 was considered statistically significant. Post-hoc multiple comparisons were performed in SciPy using Sidak’s, Tukey’s or Dunnett’s multiple comparison test as appropriate. All scripts and analyses conducted in Python were executed in a Jupyter Lab environment to ensure reproducibility.

## 3.0 Results

### 3.1 Effect of BEE on in-vitro grown Arabidopsis under abiotic stress

#### 3.1.1 Osmotic stress

In a first series of experiments, osmotic stress mitigation was tested by adding BEE at final concentration of 0.36 g/L and 0.71 g/L to plants grown in the presence of 100 and 150 mM sorbitol. In line with previous observations (Ogunsanya et al., 2022), the primary root length (PRL) significantly increased with BEE treatment in non-stressed plants compared to the control (p = 0.017 - 0.020). The application of 150 mM sorbitol inhibited primary root growth, an effect that was partially restored by adding 0.71 g/L BEE (Fig. 1A; P=0.021). A similar trend was observed for plants treated with 100mM sorbitol, albeit that this was not significant (Fig 1A; p = 0.249). The lateral root (LR) number was lower in sorbitol treated plant compared to the control (Fig 1B). A 0.71 g/L BEE application significantly increased lateral root formation under mild (100 mM sorbitol; p = 0.045) osmotic stress, which was not observed under high (150 mM sorbitol; p = 0.072) osmotic stress. The application of 0.36 g/L and 0.71 g/L BEE strongly stimulated shoot growth (Fig. 1C). Shoot growth was strongly inhibited by sorbitol treatments which was not significantly restored by BEE treatments (Fig. 1C; p = 0.245 – 0.706). These experiments showed that BEE is not very effective in protecting plants from osmotic stress.

**Figure 1.**
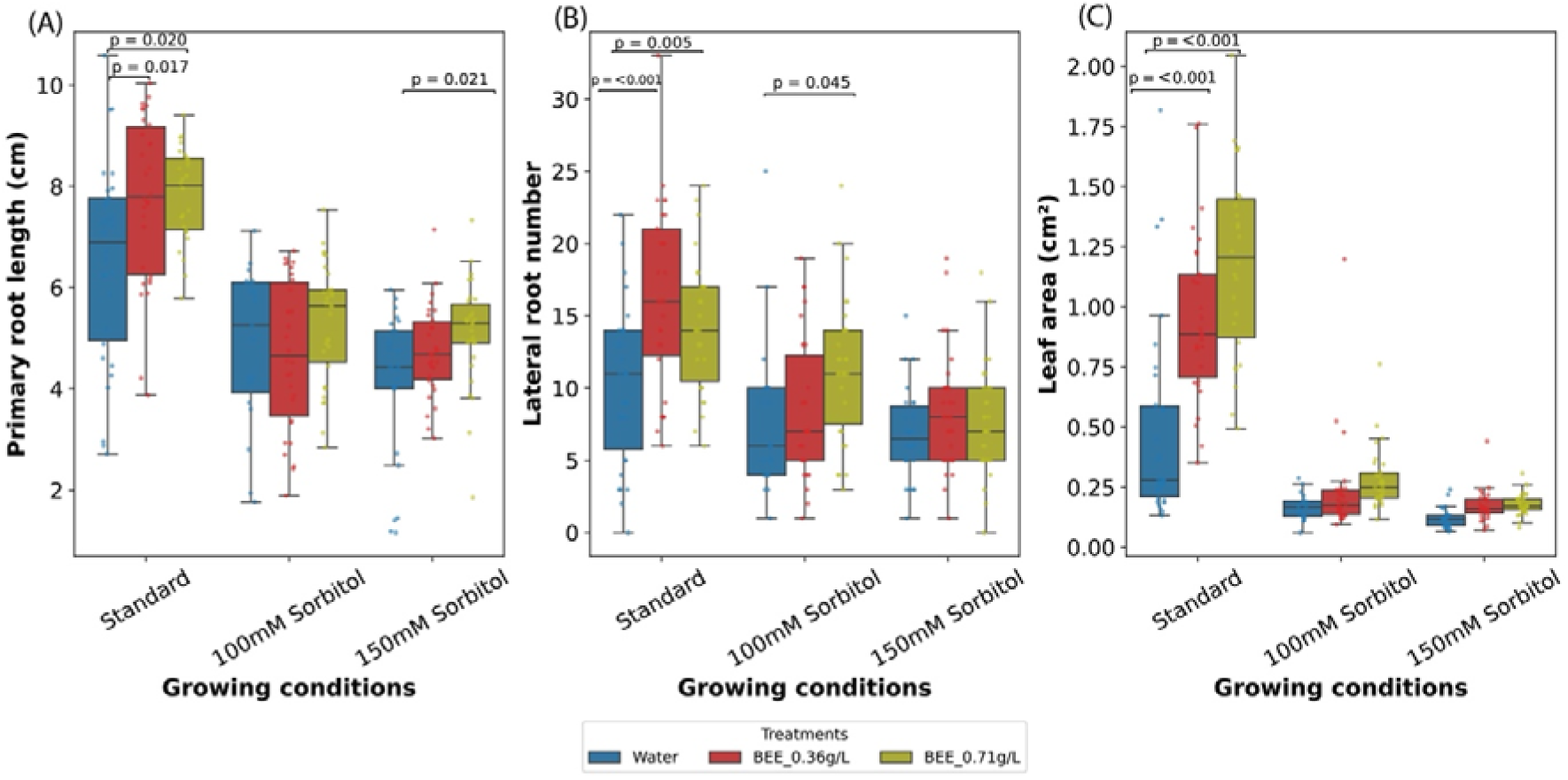
Effect of BEE on sorbitol-treated Arabidopsis plants on root and shoot growth. (A) primary root length. (B) lateral root number. (C) projected leaf area. Data represent the average of three biological and ten technical replicates per bar (30 seedlings in total, 10 per replicate). P-values indicate significant differences (p<0.05) between control (Water) and treatments (0.36g/L or 0.71g/L of BEE) in respective groups (growing condition) according to Sidak post-hoc test.

In a second set of experiments, the capacity of BEE to promote the recovering from osmotic stress was analysed. Arabidopsis plants subjected to 150 mM sorbitol for 7 days were transferred to sorbitol-free medium supplemented with 0.36 g/L or 0.71 g/L BEE. Rescued plants showed significant recovery in primary root length and lateral root number compared with non-rescued plants. Leaf area was likewise greater in rescued plants than in non-rescued controls (Fig. S1). BEE supplementation did not produce significant differences in any of these phenotypes compared with the control medium.

Overall, BEE enhanced root elongation, leaf area, and lateral root development under standard conditions, but somewhat loses its effectiveness under osmotic stress (Fig. S2).

#### 3.1.2 Salinity stress

To examine salinity stress, Arabidopsis was cultured in the presence of 100mM NaCl that reduced germination from 100±0.0% to 98.67±11.5%. BEE treatment restored germination to 99.33 ±8.2% (Fig. 2A). A Chi-square test showed no significant association between BEE treatment and germination rate under both growing conditions (χ² = 3.685 df = 3, p = 0.298), with germination ranging from 148-150 out of 150 seeds across treatments. At day 6 after sowing, the percentage true leaf was calculated as the number of emerged first “true” leaf per lot of germinated seeds. In standard conditions, 100% of BEE-treated seedlings developed true leaves, compared to 93.33 ±25.0% in control medium. The growth stimulation effect was more pronounced in 100 mM NaCl, with 61.33 ±48.9% of BEE-treated seedlings developing true leaves, compared to only 11.33 ±31.8% in the control. At 10 days after germination, leaf area was measured to evaluate vegetative growth (Fig. 2C). Under standard conditions, BEE-treated plants had a mean leaf area of 16.49 ±5.6 mm, significantly larger than the control (7.74 ±4.1mm). Similarly, under 100 mM NaCl, leaf area was reduced across all treatments, however BEE-treated plants showed a significantly larger mean leaf area (4.23 ±2.3 mm) as compared to the control (1.90 ±0.8 mm). The effect of BEE on leaf area is larger in non-stressed plants, and this is likely due to their higher baseline performance compared to stressed plants. When expressed as a relative change (ratio of +BEE/-BEE), the effect of BEE was very similar in both conditions (2.23 vs 2.13; Fig. S3), indicating that its proportional impact on leaf area is consistent irrespective of the stress status. In contrast, the +BEE/-BEE ratio for the emergence of first true leaf was highly elevated under salt stress (5.50; Fig. S3) but absent under standard condition (1.07). Together, the results shows that BEE treatment enhances Arabidopsis seedling growth under both standard and mild NaCl salt stress conditions.

**Figure 2.**
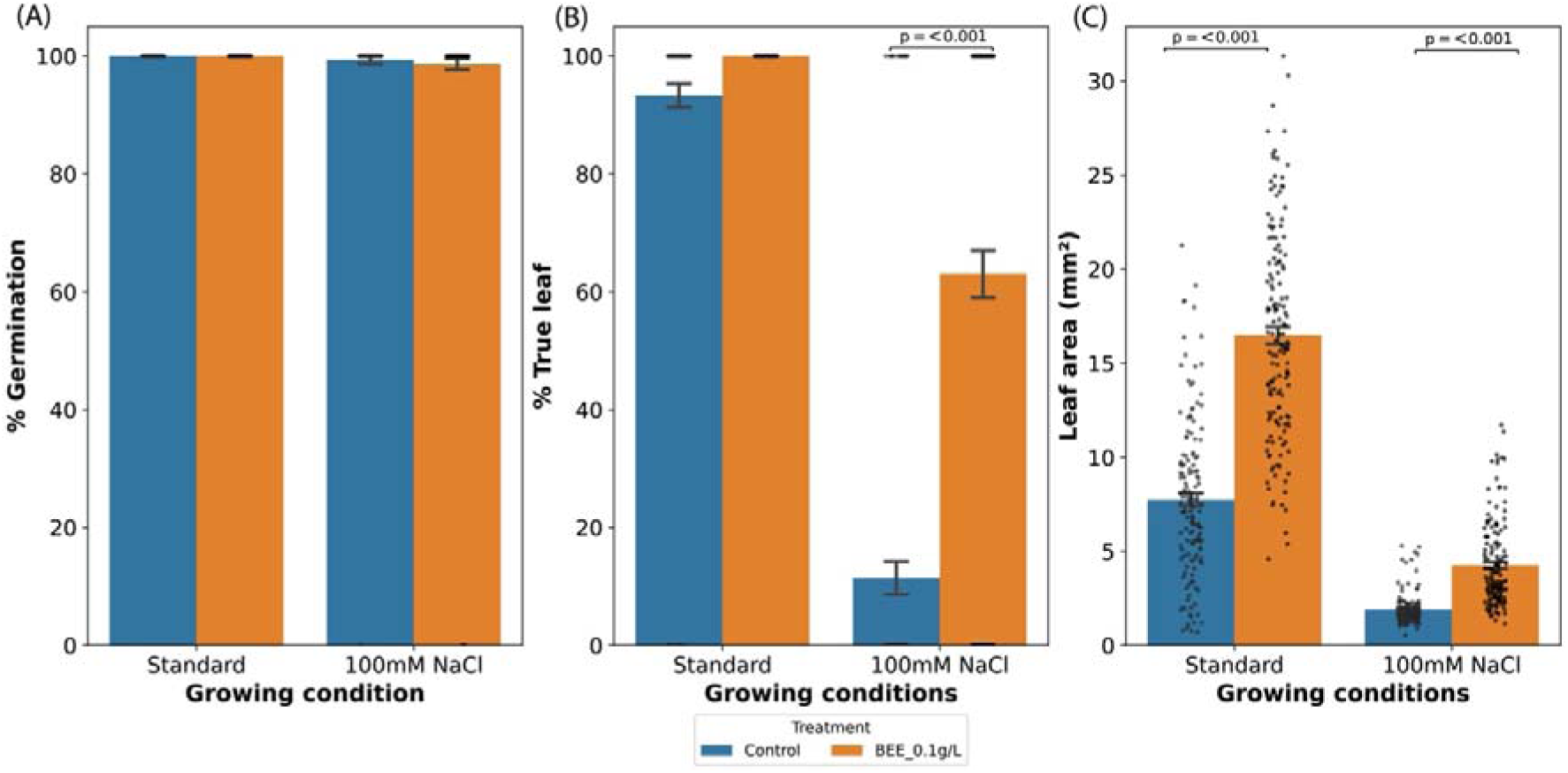
Growth parameters of Arabidopsis treated with BEE under standard and 100 mM NaCl growing condition. Shown are the effect on germination rate (A), true leaf emergence at 6DAS (B), and leaf area at 10DAS (C). P-values indicate significant differences (p<0.05, n=150) between control and BEE treatment in respective groups (growing condition) according to TukeyHSD test.

The effect of BEE on Arabidopsis shoot health was evaluated by measuring chlorophyll fluorescence (Fv/Fm), chlorophyll index (ChlIdx), and anthocyanin accumulation (mARI). BEE-treated plants showed a Fv/Fm ratio slightly higher than control (0.67 ±0.07, and 0.71 ±0.05 respectively). Under mild salt stress, BEE treatment sustained a higher ratio Fv/Fm compared to the control (0.65 ±0.04 vs. 0.56 ±0.06 Fig. 3A). The chlorophyll index, a quantification of leaf greenness, showed that BEE-treated plants contained a significantly higher chlorophyll content under both standard and salt stress conditions (Fig. 3B). Anthocyanins levels were significantly higher in BEE-treated plants. (Fig. 3C). Anthocyanins are photo-protectants enhancing chlorophyll fluorescence, chlorophyll index, and leaf expansion (Gould, 2004; Steyn et al., 2002), suggesting that BEE-treatment contributes to stress alleviation.

**Figure 3.**
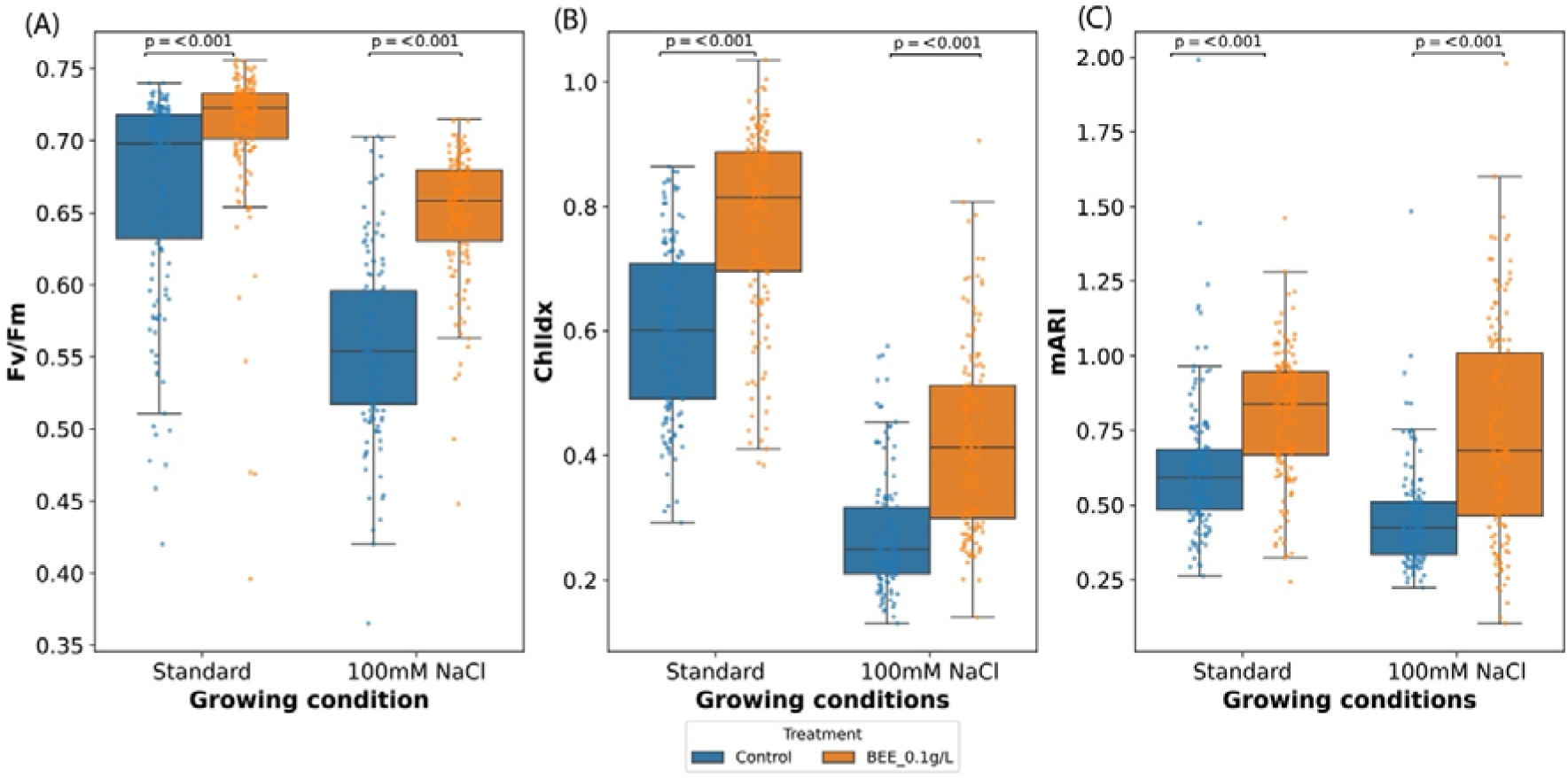
Health parameters of Arabidopsis treated with BEE under standard and 100 mM NaCl growing condition. Effect on chlorophyll fluorescence (A), chlorophyll index (B), and modified anthocyanin reflective index (C). Data represent the average of three biological and fifty technical replicates per bar (150 seedlings in total, 50 per replicate). Statistical significance (n=150, p<0.05) between control and treatment BEE in respective groups (growing condition) was performed using Two-way ANOVA, followed by TukeyHSD test.

### 3.1 Effect of BEE on Arabidopsis grown on substrate

#### 3.1.1 Vegetative growth

In a next series of experiments, we evaluated whether the growth stimulation effect of BEE on *in vitro* cultured Arabidopsis is reproduced in soil conditions. Arabidopsis seedlings of 10 days old were either drenched (root treatment) or sprayed (shoot treatment) with 0.18 g/L and 0.36 g/L BEE. To allow comparison with the *in vitro* test conditions, it required autoclaving of BEE, (Auto_BEE) and we included untreated (not autoclaved; NA_BEE) BEE as a reference. At 0.18 g/L, both BEE solutions did not alter the leaf area, suggesting that shoot growth stimulation as observed in the *in-vitro* experiments was absent (Fig 4). The 0.36 g/L autoclaved BEE solution applied by root drenching significantly increased the shoot area whereas spraying the same solution did not exert the growth stimulation (Fig. 4). These observations support the biostimulant effect of BEE and suggest that its activity increases by autoclaving and that it is active through the root system.

**Figure 4.**
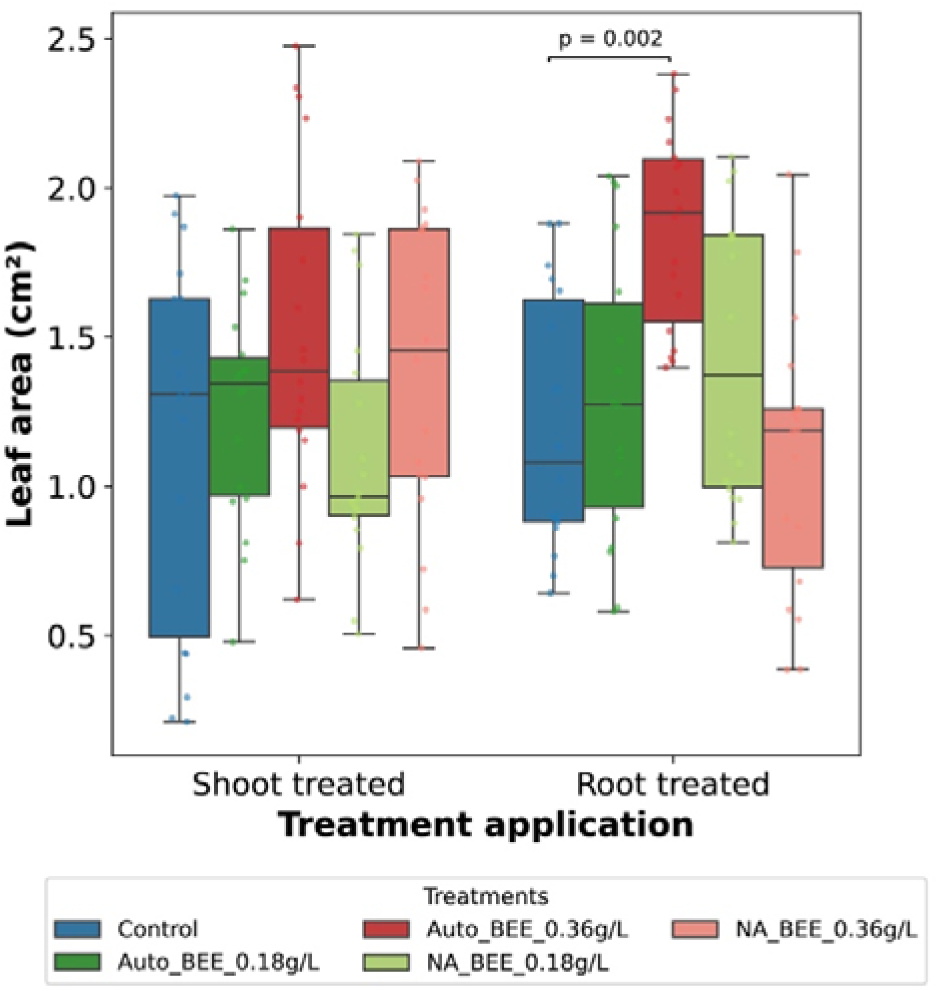
Effect of different forms of BEE and different application on Arabidopsis leaf area. Data represent eighteen technical replicates per bar. Graph is presented in a boxplot with all datapoints and bars indicating minimum and maximum value. Statistical significance (n=18, p<0.05) between control and treatment BEE in respective groups (application) are shown and performed using Two-way ANOVA, followed by TukeyHSD test. Auto_BEE: autoclaved Belgian endive extract, NA_BEE: non-autoclaved Belgian endive extract.

#### 3.1.2 Lettuce cultivation with BEE

Lettuce seedlings were treated with 0.18 g/L or 0.36 g/L BEE, grown for 31 days after BEE treatment (DAT), after which the mean fresh shoot weight was compared with that of untreated plants. The BEE-treated plants did not show a significant difference in leaf fresh weight (Fig. S4A) or dry weight (Fig. S4B) from the untreated control. The NDVI optical properties of lettuce leaf samples was higher in plants treated with 0.18 g/L BEE (P=0.021) compared to the control but the higher BEE dose (0.36 g/L) did not show a significant effect (Fig. 5A). The leaf greenness (GM1 index) was higher (P=0.029, Fig. 5B) at 0.18 g/L BEE compared to control and the 0.36 g/L BEE treatment, and the carotenoid content (CRI1 index) was also higher after 0.18 g/L BEE treatment (P=0.002, Fig. 5C) but not with 0.36 g/L BEE. No significant differences were observed in anthocyanin content (ARI1), general coefficient (G), and the internal reference (Ctr1).

**Figure 5.**
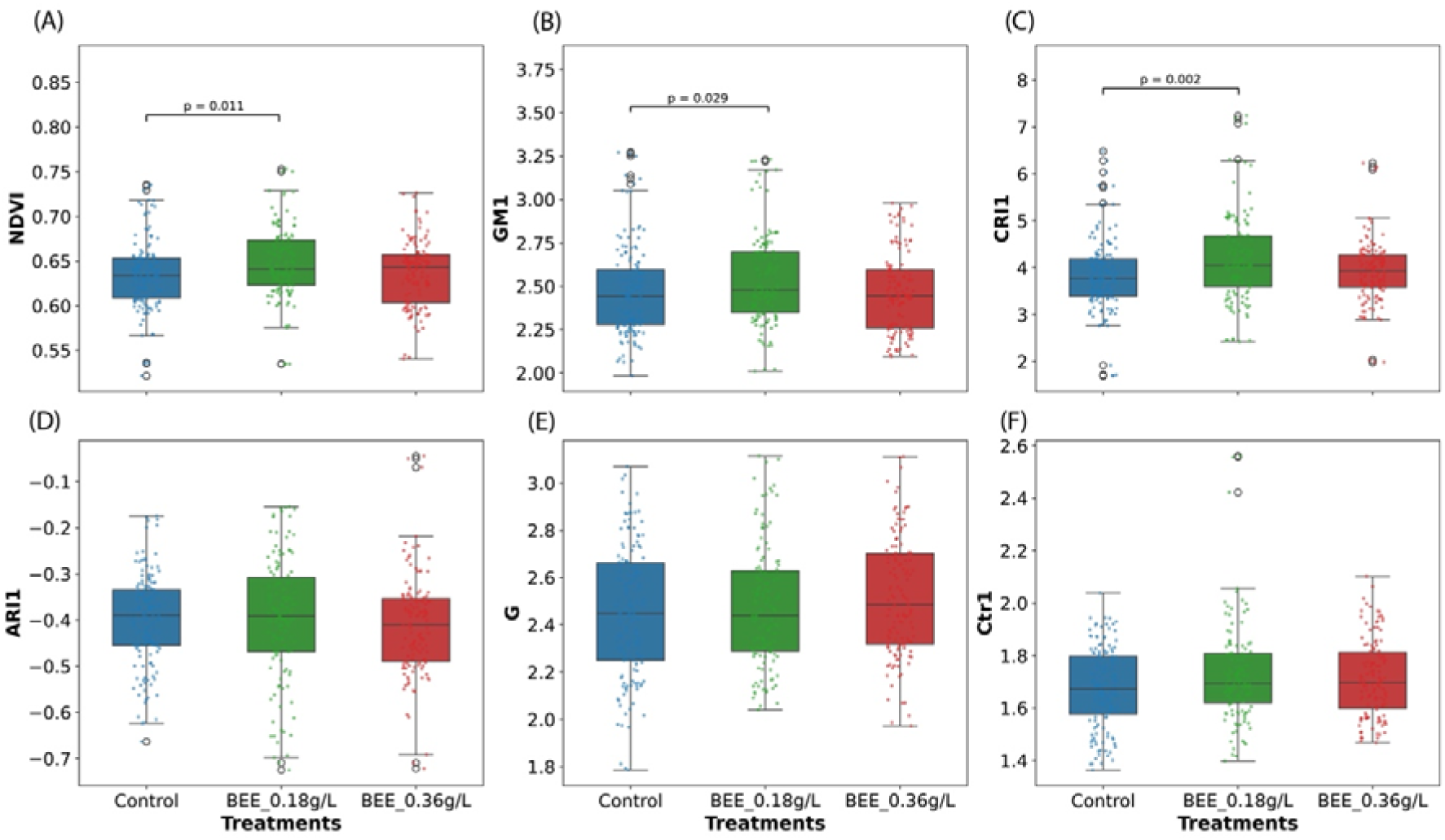
Graphical representation of the effect of BEE treatment or not (Control) on Lettuce leaf optical indices. (A) NDVI: Normalized Difference Vegetation Index. (B) GM1: Greenness Model 1. (C) CRI1: Carotenoid Reflectance Index 1. (D) ARI1: Anthocyanin Reflectance Index 1. (E) G: General coefficient. (F) Ctr1: Control Value 1. P-values <0.05 according to Dunnett’s multiple comparison test preceded by one-way ANOVA are displayed on the graphs.

### 3.2 Effect of BEE on sweet pepper growth and fruit development

#### 3.2.1 Impact of BEE on sweet pepper grown in a greenhouse

Sweet pepper experiments were conducted in a greenhouse over the course of three consecutive years. In a first round of tests (year 2022), the sweet pepper plants treated with BEE were similar in size compared to the control group, reaching an average height of 113.2 ± 4.1 cm, and 103.6 ± 12.4 cm respectively at the end of the experiment (Fig. 6A). DUALEX measurements indicated that the chlorophyll content in BEE-treated plants was significantly higher (40.0 ± 1.2 µg/cm^2^, p = 0.009) compared to control plants (37.2 ±1.3 µg/cm^2^; p ≤ 0.05, Fig. 6B). The result suggests that BEE treated plants possess higher photosynthetic activity which may lead to higher capacity of carbon dioxide fixation.

**Figure 6.**
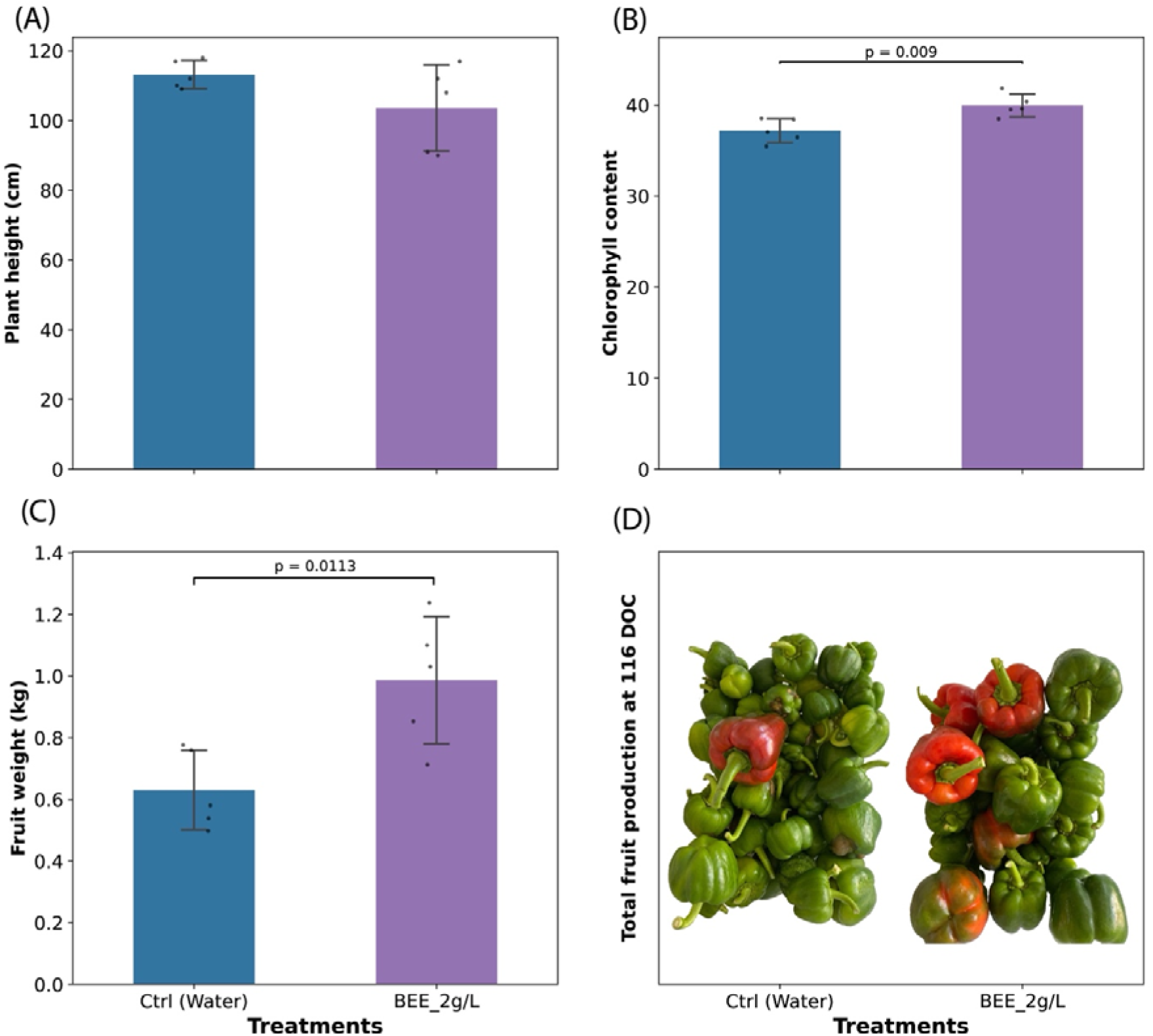
Effect of BEE treatment or not on growth parameters and yield of sweet pepper plants under greenhouse conditions. (A) The average plant height. (n=5). (B) The chlorophyll content from DUALEX readings. (n=5). (C) The total fresh weight of all fruits per treatment. (n=5). (D) Pictures comparing the mature fruits from treated and untreated groups. All graphs are presented with error bars indicating standard deviation, and p-values of statistical significance according to t-test. BEE: Belgian endive extract. Ctrl: Control treated with water.

Pepper plants treated with BEE showed a 56% increase in yield (0.99 ± 0.2 kg of fruits per plant) compared to the control with (0.63 ± 0.1 kg of fruits per plant) at 116 days of cultivation (Fig. 6C). In addition, BEE-treated plants produced fruit that matured earlier than that of the untreated control (Fig 6D).

#### 3.2.2 Impact of BEE on sweet pepper grown indoor under artificial light

To test whether the effect of BEE on sweet pepper fruit development and maturation could be replicated under artificial light condition, plants were cultured in a controlled environment; plant growth room (year 2023). In this trial, young sweet pepper plants were transplanted onto rockwool slabs and fruit production was evaluated 97 days after cultivation, by counting the number of fruits per plant from treated and non-treated plants. The average number of fruits per plant was 11.3 (± 3.6) per plant in the BEE-treated group whereas the control group had 12.8 (± 6.1) fruits per plant (Fig. 7A). A significant effect was observed for fruit yield with BEE producing on average of 0.29 (± 0.2 kg) fruits per plant and 0.13 (±0.1) kg fruits per plant in the control group (p = 0.043).

**Figure 7.**
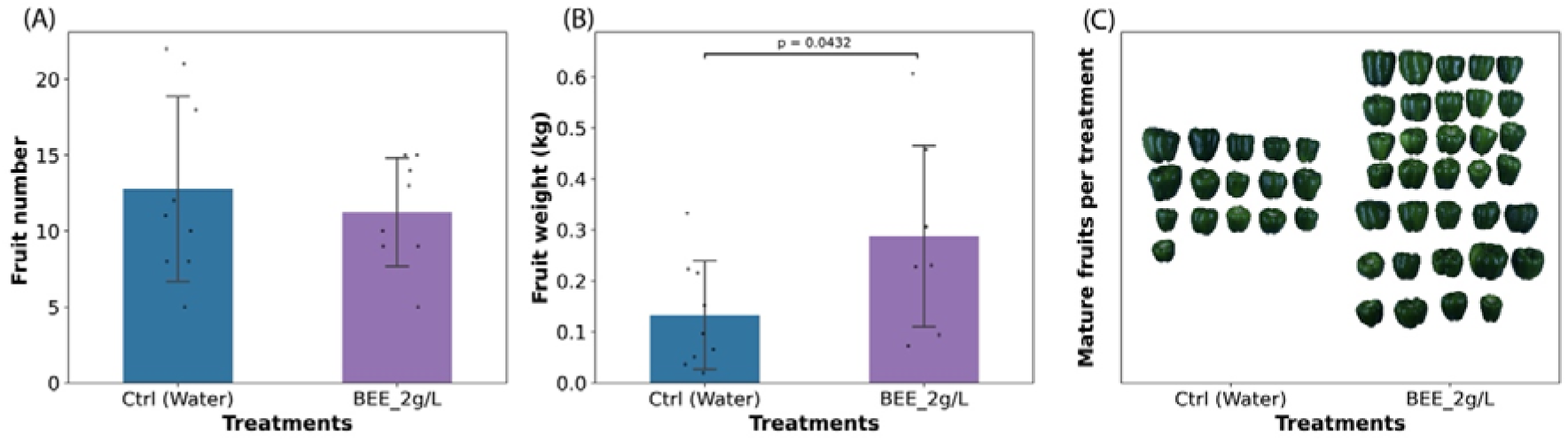
Effect of BEE treatment or not on yield of sweet pepper plants under growth room conditions. (A) The average fruit number. (n=8-9). (B) The total fresh weight of all fruits per treatment. (n=8-9, p ≤ 0.05). (C) Pictures comparing the mature fruits from treated and untreated groups. All graphs are presented with error bars indicating standard deviation, and statistical significance (indicated by p-values) was determined using t-test. BEE: Belgian endive extract. Ctrl: Control treated with water.

More fruit biomass was produced upon BEE treatment (Fig 7B). Although the number of fruits per plant in the control group is more than the number of fruits per plant in the BEE-treated group, the number of mature fruits in the BEE-treated group was twice that of the mature fruits in the control group (Fig 7C). These results further support the growth promoting effect of BEE on sweet pepper fruit yield.

### 3.3 The fruit development effect of BEE

The third pepper experiment (year 2024) included 30 plants that were grown in a greenhouse, and fruits were analysed at 116 days of cultivation. The lower branches of the plants were pruned before flowering to promote fruit ripening. As in the previous experiments, BEE treatment consistently enhanced fruit development producing larger fruit that ripened earlier. As a measure for yield, we determined the number of red (ripe) fruits and the distribution of mature fruits per plant (Fig. 8). The BEE-treated plants produced more red fruits (0.47 ± 0.6) than the control (0.06 ± 0.2) consistent with the effect of BEE on paprika fruit ripeness and yield in previous experiments. A chi-square test showed a significant difference in the distribution of red fruits between groups (χ² = 6.76, df = 2, p = 0.034). This difference was mainly driven by a higher proportion of plants with ≥1 red fruits in the treatment group and a higher proportion of plants with zero red fruits in the control group. However, a presumed earlier maturation induced by BEE was not reflected in a significant difference in plant height, chlorophyll content, fruit number and fruit weight (Figs. S5A-D).

**Figure 8.**
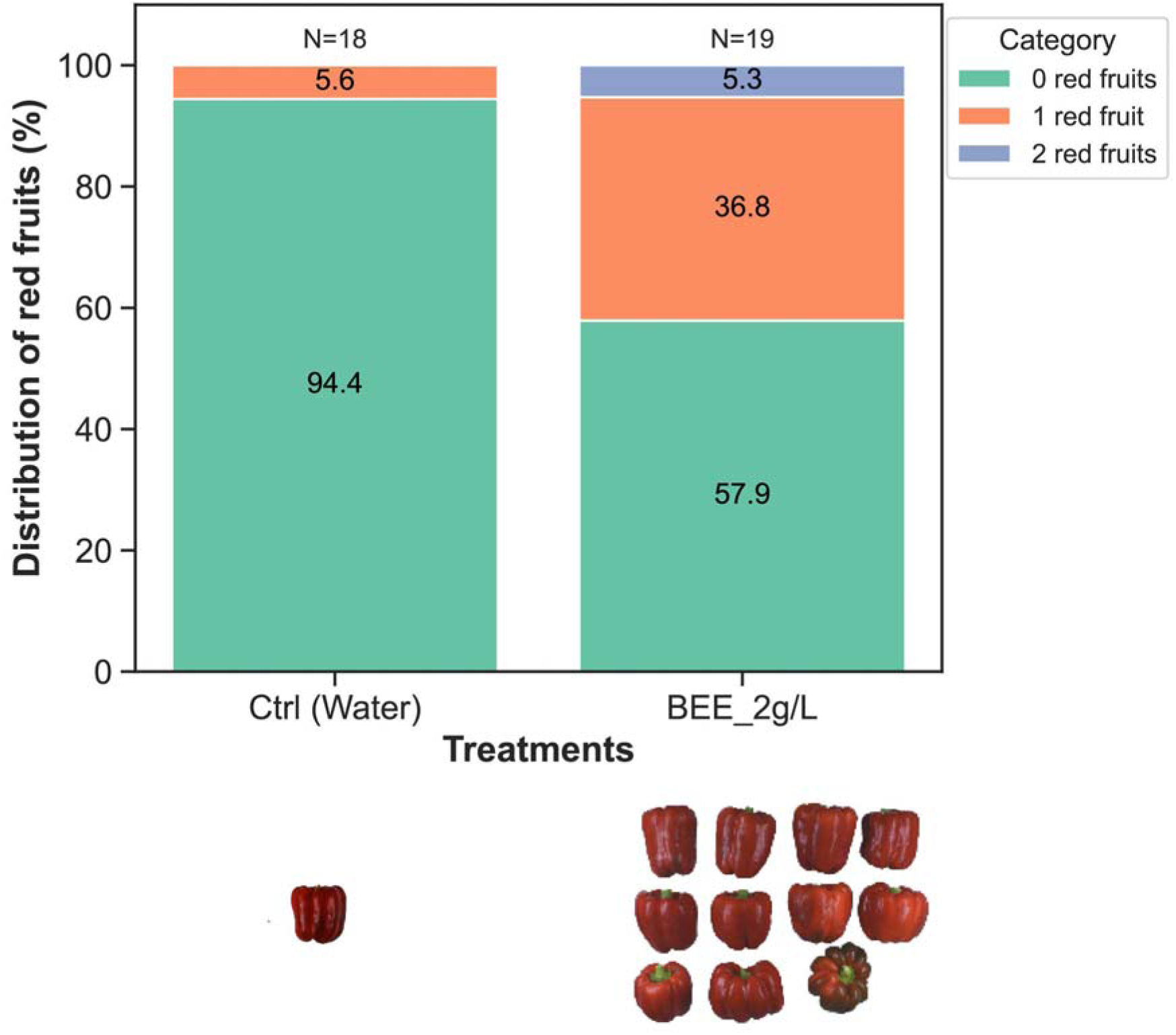
Effect of BEE treatment or not on maturity and ripeness of sweet pepper fruits under greenhouse conditions. Stacked graph of the percentage distribution of ripe (red) fruits per plant per treatment (top). Pictures comparing the ripe (red) fruits from treated and untreated groups (bottom). BEE: Belgian endive extract. Ctrl: Control treated with water.

As sweet pepper fruit ripening is associated with ethylene signalling (Huang et al., 2024) ethylene levels were analysed in young leaves of 54-days-old plants. BEE-treated plants released a similar level of ethylene as non-treated plants 48 hours post-treatment (Fig. S6). This result suggests that the previously observed accelerated ripening effect in sweet pepper fruits may not be directly linked to a change in ethylene levels.

### 3.4 Effect of BEE treatment on sweet pepper under drought stress

Greenhouse pepper cultures (Year 2024) were subjected to drought stress by interrupting irrigation from day 56 to day 60 when the soil moisture reached <30%. BEE was applied on day 61. Height, chlorophyll content, fruit weight, and fruit number were not affected by BEE treatment (Fig. 9A-D). However, the number of ripe fruits (fully red coloured), was much higher in the treated plants, resulting in a higher mean percentage of ripened fruits in BEE-treated (30.85%) plants compared to the untreated control (16.95%), albeit not significant. The chi-square test did not show a statistically significant difference in the distribution of red fruits between groups (χ² = 3.47, df = 2, p = 0.176). However, the observed distributions suggest a trend toward higher frequencies of plants with two red fruits in the treatment group (Fig. 9E). A similar trend was observed in another experiment (Year 2022) where BEE treatment only increased fruit maturity (Fig. S7). These findings suggest that while BEE may not directly enhance vegetative growth under drought stress, it tends to promote fruit maturity and ripeness.

**Figure 9.**
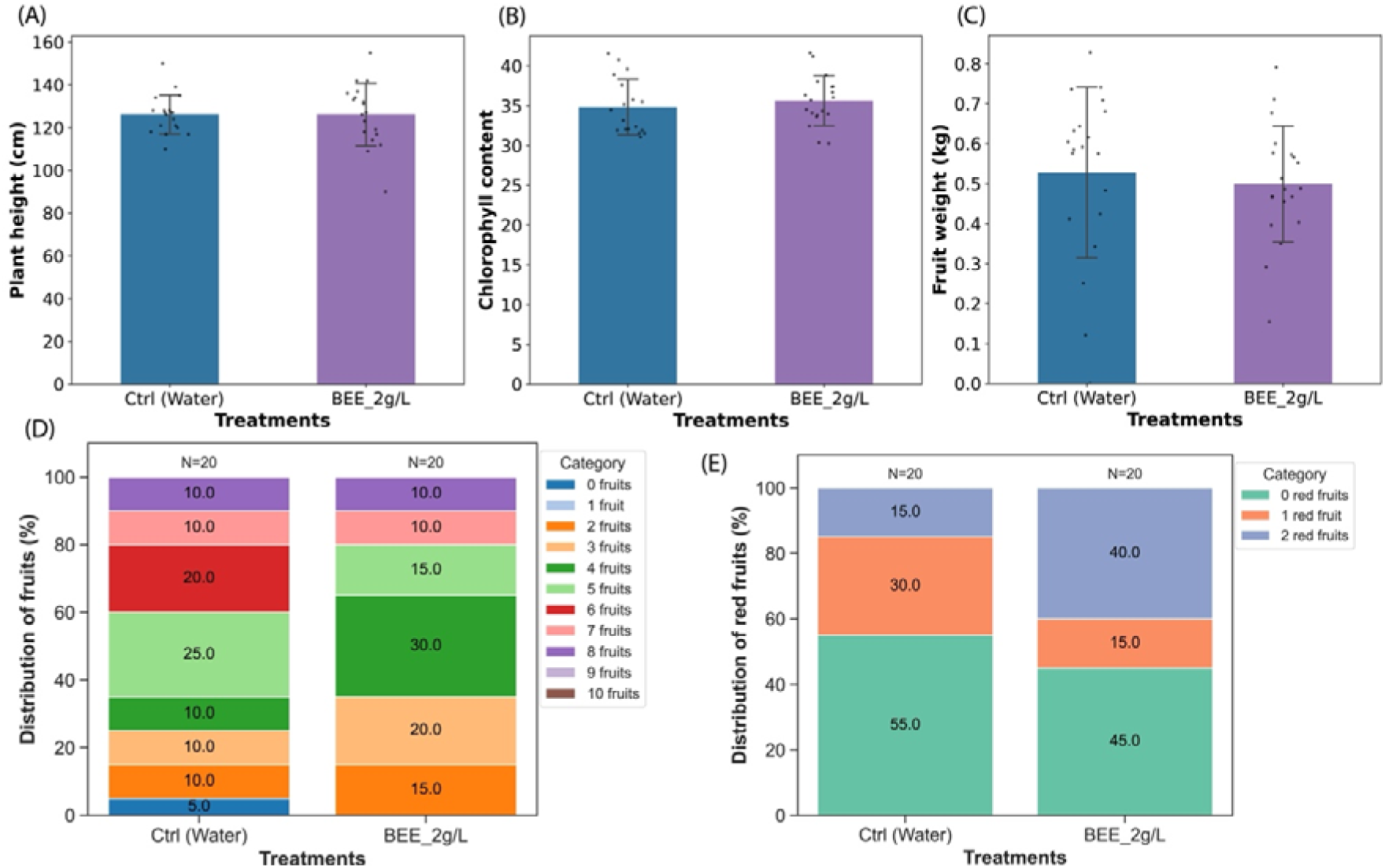
Effect of BEE treatment or not on growth parameters and yield of drought stressed sweet pepper plants under greenhouse conditions in 2024. (A) The average plant height (n=18-19). (B) The chlorophyll content from DUALEX readings (n=18-19). (C) The total fresh weight of all fruits per treatment (n=18-19). (D) The percentage distribution of fruit number per treatment. (E) Percentage distribution of the number of ripe (red) fruits per plant per treatment. Graphs are presented with error bars indicating standard deviation and statistical significance was performed using t-test.

## 4.0 Discussion

In this study, we set out a series of experiments to investigate the potential beneficial activity of BEE under conditions of stress and whether it can be used in the context of crop production, *in casu* lettuce and sweet pepper.

### 4.1 Effect of BEE on abiotic stress

Salinity stress negatively impacts plant growth by inhibiting seed germination, stunting root and shoot development, and ultimately reducing yield. Biostimulants mitigate these effects by promoting root growth and biomass accumulation (Alzate Zuluaga et al., 2022; Ntanasi et al., 2024; Zuzunaga-Rosas et al., 2023). Plant extract-derived biostimulants such as protein hydrolysates have been effective in improving biomass production under salinity stress in crops like maize (Colla et al., 2014; Ertani et al., 2013). In this study, the biostimulant BEE had no effect on germination under salt stress, but mitigated the stress as evidenced by early true leaf emergence, larger leaf area and improved plant health compared to untreated plants. BEE either (1) actively enhances the plant’s stress tolerance, or (2) it promotes early plant vigour such that plants are more developed and tolerate better the salt treatment. As BEE was applied prior to germination and did not affect the germination rate it appears to directly protect from salt stress and not through an early growth advantage. BEE enhanced photosynthetic efficiency, thereby contributing to increased biomass and leaf area. Since leaves serve as the primary sites of photosynthesis (Lv et al., 2023), the enhanced effect of BEE on early true leaf emergence under stress may confer an advantage to the photosynthetic performance of the stressed plants, giving them an edge over untreated control. This improvement may, in turn, promote larger leaf area development, creating a positive feedback cycle that further enhances growth under stress conditions.

Salinity stress affects photosynthesis by accelerating chlorophyll degradation and inducing stomatal closure, which limits CO_2_ intake. Biostimulant application has been reported to reduce chlorophyll degradation while enhancing nutrient uptake and maintaining plant productivity (Ikuyinminu et al., 2023; Zuzunaga-Rosas et al., 2022). BEE treatment improved chlorophyll fluorescence and chlorophyll index, contributing to overall plant health under salt stress. Under standard conditions, the increased anthocyanin accumulation in BEE-treated plants is unlikely to signal oxidative stress, as overall plant health and growth parameters remained unaffected. However, mARI may still function as a photo-protectant, particularly in young plants, as Fv/Fm values were only slightly above the 0.7 threshold, suggesting that the spectral measurements might have detected a mild stress response in developing seedlings. Concisely, BEE treatment enhances Arabidopsis health by improving photosynthetic efficiency, chlorophyll content, and anthocyanin accumulation, particularly under mild salt stress. The higher Fv/Fm values, chlorophyll index, and mARI suggest BEE mitigates stress effects while potentially acting as a photo-protectant in young seedlings. These findings highlight BEE’s potential in enhancing plant resilience and salinity stress adaptation. Additional benefits of biostimulants in mitigating salinity stress include enhanced antioxidant activity (Li, Evon, et al., 2022; Naboulsi et al., 2022) and improved ion homeostasis and osmotic regulation (D’amato & Del Buono, 2021). The results from this study align with the established role of biostimulants in alleviating salt stress.

In addition to salt toxicity, we analysed osmotic (drought) stress by transferring seedlings to medium containing 100 mM and 150 mM sorbitol. There was no significant alleviation of osmotic stress under *in vitro* conditions, nor did we observe an advantage of BEE treatment with plants subjected to drought stress. Similarly, *ex vitro* experiments confirmed that BEE did not enhance growth parameters in drought-stressed sweet pepper plants compared to controls. These findings contrast with previous studies reporting that biostimulants mitigate drought stress by improving growth and yield (De Clercq et al., 2023) enhancing photosynthetic efficiency (Soussani et al., 2023), and optimizing nutrient and water use efficiency (Repke et al., 2022).

Overall, these results suggest that BEE does not mitigate abiotic stress by reducing osmotic pressure, but rather by improving photosynthetic efficiency and biomass accumulation. The potential impact of BEE on metabolic processes and stress-related signalling pathways was not addressed in this study.

### 4.2 Evaluation of BEE bioactivity ex vitro

The use of both model plants and economically important crops in biostimulant research is essential for bridging fundamental and applied studies. Model plants, such as *Arabidopsis thaliana*, provide critical insights into molecular pathways, gene expression, and physiological responses triggered by biostimulants. In contrast, crops and horticultural plants validate these findings under field conditions, ensuring practical applicability for large-scale agriculture (Rouphael & Colla, 2020). This dual approach strengthens the scientific foundation of biostimulant research while promoting sustainable agricultural practices. In *in vitro* experiments, BEE treatment significantly increased leaf area in Arabidopsis, serving as a key indicator of shoot growth. Given this significant growth enhancement, it was necessary to evaluate whether these effects could be replicated under *ex vitro* conditions in both Arabidopsis and crop plants. Initial *ex vitro* experiments on Arabidopsis tested two forms of BEE, as *in vitro* applications typically involve autoclaving BEE with the growth medium. When BEE was not autoclaved in an *in vitro* experiment, some treated plants failed to outperform controls, while others showed contamination (Supplementary Fig. S8). This suggests that heat treatment enhances the bioactivity of BEE, with heat-treated BEE demonstrating the most significant effects on both vegetative and reproductive growth.

Translating *in vitro* findings to *ex vitro* conditions also requires the understanding of how plants absorb treatments. In *in vitro* experiments, BEE is uniformly available to both roots and shoots tissues in contact with the medium, whereas in *ex vitro* conditions, uptake depends on the application method with clear biostimulant effects observed after root drenching and now impact from shoot spraying. Root drenching typically enhances root development and nutrient uptake (Colla et al., 2015; Lucini et al., 2015, 2018), while foliar application targets photosynthesis, metabolism, and stress responses (Cristiano et al., 2018; Lucini et al., 2015; Peroni et al., 2024). Although root drenching was the most effective method for BEE application, a combination of root and foliar application may provide additional benefits, as seen in *in vitro* setup and suggested by Lucini et al., (Lucini et al., 2015).

The initial *ex vitro* results revealed insight on the application strategy of BEE in crops. In the lettuce experiment, the heat-treated BEE was applied via root drenching. No significant difference was observed in shoot fresh and dry weight between the BEE-treated and control plants. Leaf area, measured in other experiments, could not be assessed here due to the characteristics of the Expertise variety, hence shoot weight. The comparable differences in the shoot fresh and dry weights indicates that the similarity in the fresh weight was not due to differences in water retention, suggesting that BEE did not influence leaf water retention. BEE treatment, however, promoted plant greenness and health, as evidenced by significantly higher leaf indices (NDVI and GM1) associated with photosynthetic activity and general plant health. This is consistent with the previous results of BEE enhancing plant performance and health under salt stress. In addition, BEE increased the carotenoid content.

Previous studies have shown that plant-derived biostimulants increased leaf biomass in the short term and enhance reproductive traits, including fruit number and weight in *Capsicum* spp. in the long term (Ertani et al., 2014). While cultivation practices differed across the three pepper experiments, the decline in efficacy of BEE in the third year is likely attributable to the long-term storage of BEE and associated slow decay in bioactivity. Variability in performance has been reported for some biostimulants. For instance, Ertani et al. (2015), reported that grape skin extract and alfalfa hydrolysates both increased fruit quality in chili pepper, yet differed in effectiveness due to their origin, processing methods, and endogenous hormone content such as indole-3-acetic acid (Ertani et al., 2015). More broadly, different groups of biostimulants, including plant and algae extract, protein hydrolysates, and humic substances, have been shown to enhance fruit traits and quality such as antioxidant capacity, ascorbic acid, carotenoids, and capsaicinoids across different crops (Rodrigues et al., 2020). These studies emphasize that biostimulant responses are product-0 and cultivar-specific, necessitating targeted optimization strategies.

A notable consideration in interpreting biostimulant effects is the time delay between application and phenotypic response. In many cases, this lag phase reflects indirect mechanisms, such as modulation of soil-plant-microbe interaction and gradual shifts in nutrient availability and signalling pathways (Hellequin et al., 2020). Similarly, complex biomolecules such as polysaccharides e.g., in seaweed extracts require (microbial) degradation before becoming active (Andreotti et al., 2022; Hellequin et al., 2020). Additionally, when applied during early developmental stages, biostimulants often act as priming agents, influencing downstream processes such as flowering, fruit set, and ripening (Kazakov et al., 2024; Staykov et al., 2025). However, in this study this delay was likely minimized due to the activation of BEE done by heat treatment before application, which may have increased the immediate availability of active compounds. Furthermore, while studies report cumulative benefit from repeated applications (Staykov et al., 2025), the single application and one-time harvest design suggest that observed responses reflect short-term effects, although cumulative impacts cannot be excluded under repeated application scenarios.

A key trait of BEE treatment that remained consistent was the acceleration of fruit ripening. While ethylene is a central regulator of fruit ripening, no significant increase in ethylene production was detected within 48 hours of BEE application, indicating that BEE does not trigger immediate ethylene spike. Instead, the observed effect may arise from indirect modulation of developmental pathways, potentially through improved fruit set, or priming of metabolic processes that later influence ripening dynamics. This interpretation is consistent with the concept that biostimulants act through complex regulatory networks rather than single hormonal pathways.

Overall, these findings reinforce the importance of considering product stability, biochemical composition, and application timing when evaluating biostimulant performance. Future work should focus on mechanistic understanding of BEE, particularly in its interaction with hormonal signaling and developmental processes, as well as its stability during storage. Such insights will be essential for optimizing its use and ensuring consistent agronomic outcomes.

## 5.0 Conclusion

In conclusion, the application of BEE demonstrates significant potential in enhancing various aspects of plant development, including vegetative growth and fruit quality. Furthermore, the results corroborate that the growth-promoting effects of BEE observed *in vitro* can be translated to *ex vitro* cultivation systems. Notably, root application of heat-treated BEE was the most effective strategy, underscoring the importance of both application method and treatment form in maximizing biostimulant efficacy. This study contributes to the growing body of research that emphasises the role of biostimulants in promoting plant vigour, improving nutrient uptake, and increasing tolerance to abiotic stress. It particularly provides a strategy for streamlining the identification of bioactivity in newly discovered biostimulants while highlighting the importance of translating *in vitro* findings to *ex vitro* conditions.

