## Supplementary file for "Belgian endive-derived biostimulant activity in Arabidopsis, lettuce and sweet pepper at different developmental stages, environmental conditions, and application methods"

Table S1. Overview of the spectral parameters used in this study for the estimation of Arabidopsis health status

| Parameter | Formula | Physiological importance | Source |
|---|---|---|---|
| $F_0$ | | Minimal level of fluorescence measured at 730nm* after exposure to a weak measuring beam. | (Baker, 2008; Murchie & Lawson, 2013) |
| $F_m$ | | Maximum level of fluorescence measured at 730nm* after exposure to a brief saturating pulse. | (Baker, 2008; Murchie & Lawson, 2013) |
| $\frac{F_v}{F_m}$ | $\frac{F_m - F_0}{F_m}$ | Chlorophyll fluorescence; Efficiency of photosystem II in a dark-adapted state (PSII). | (Baker, 2008; Murchie & Lawson, 2013) |
| ChlIdx | $\frac{\rho_{770}}{\rho_{710}} - 1$ | Chlorophyll index; Vegetation index for the estimation of the chlorophyll content in leaves. | (Gitelson et al., 2003) |
| mARI | $\left(\frac{1}{\rho_{550nm}} - \frac{1}{\rho_{710nm}}\right) \rho_{770nm}$ | Modified Anthocyanin Reflectance Index; Gives an estimation of the anthocyanin-content in leaves. | (Gitelson et al., 2009) |

*Spectral width of 40 nm at full width half maximum.

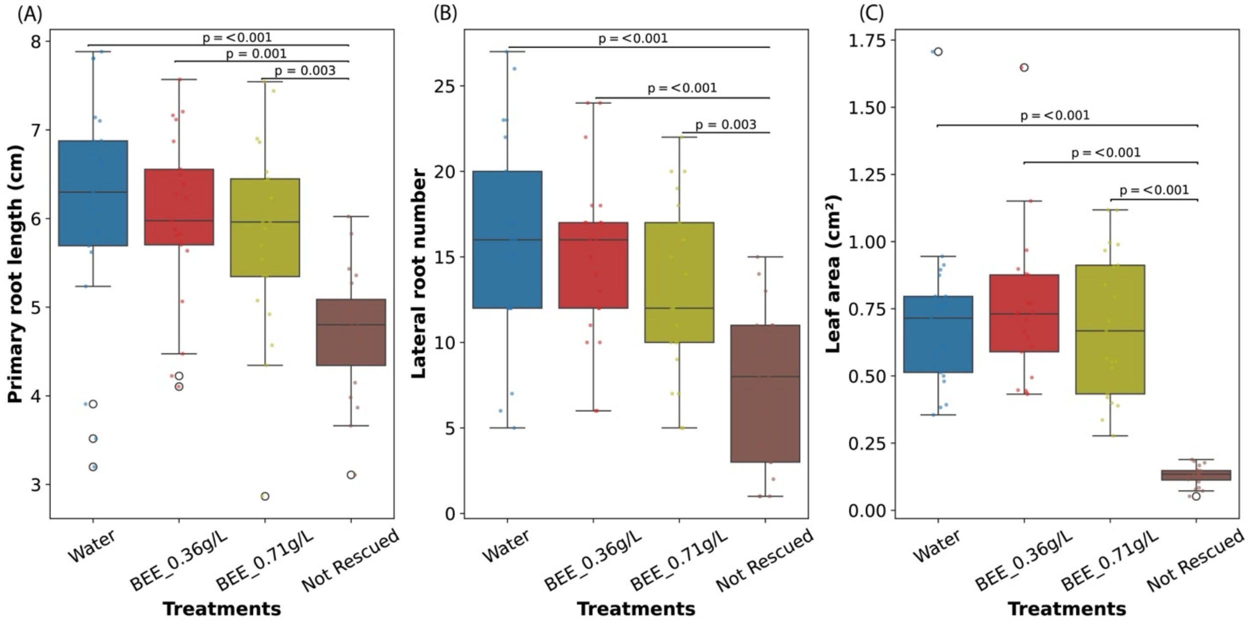


Figure S1. Graphs showing the ability of BEE to rescue osmotic stressed plants. (a) shows the primary root length of plants rescued from 150mM Sorbitol to either BEE (0.36g/Lor 0.71g/L), water (control), or not rescued. (b) and (c) show the effect of BEE on the lateral root and the rosette area respectively. Data represent the average of three biological and seven technical replicates per bar (21 seedlings in total, 7 per replicate). P-values indicate significant differences ($p<0.05$) between non-rescued and rescued plants according to Tukey's multiple comparison test.

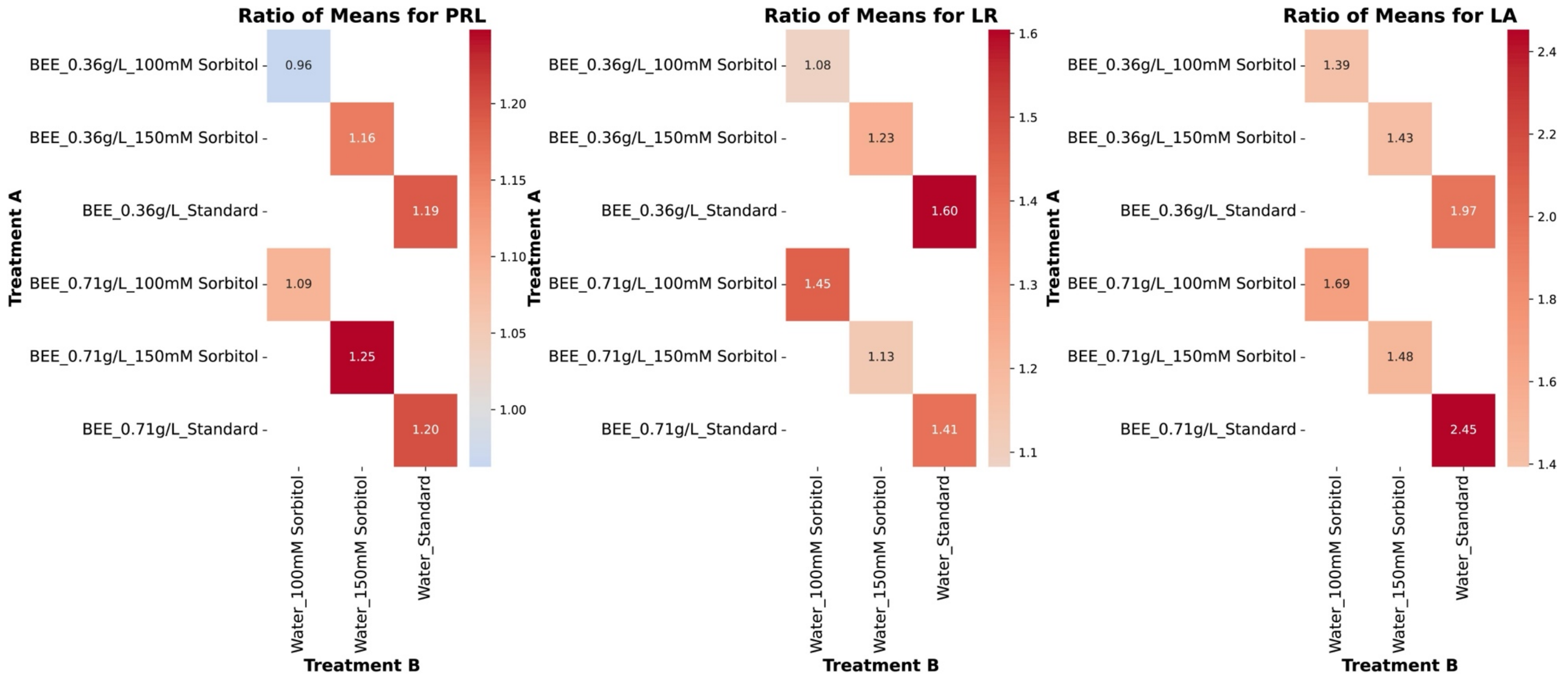


Figure S2. Ratio between BEE application or not in a sorbitol or sorbitol-free medium.

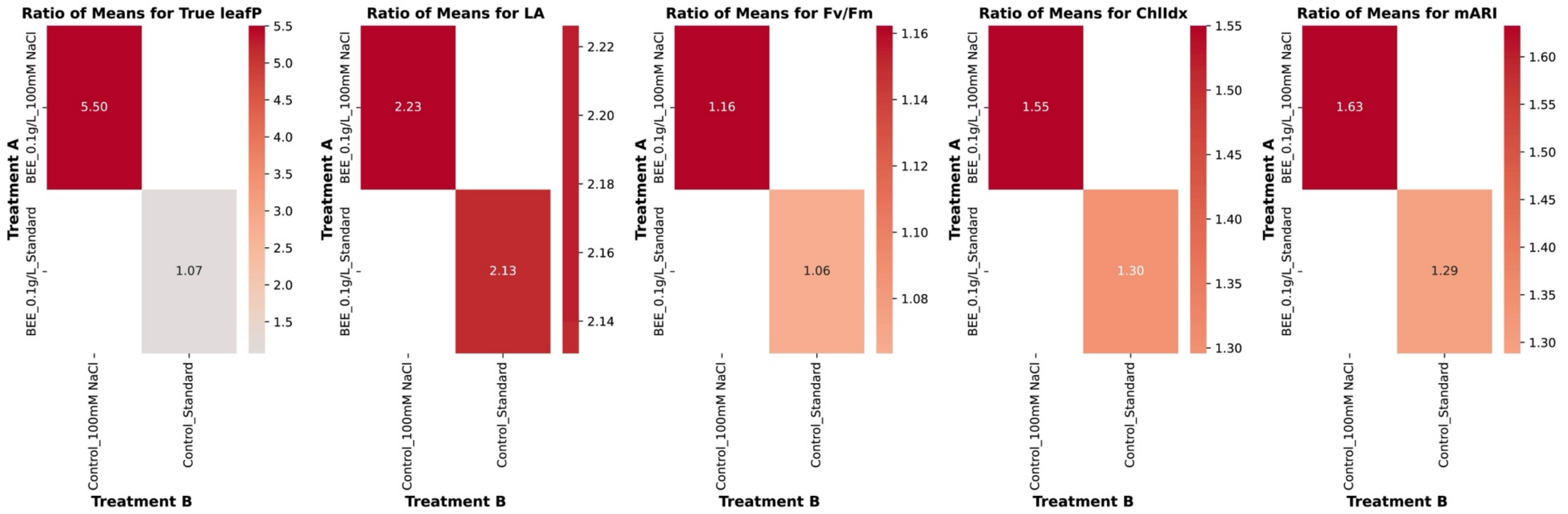


Figure S3. Ratio between BEE application or not in a NaCl or NaCl-free medium.

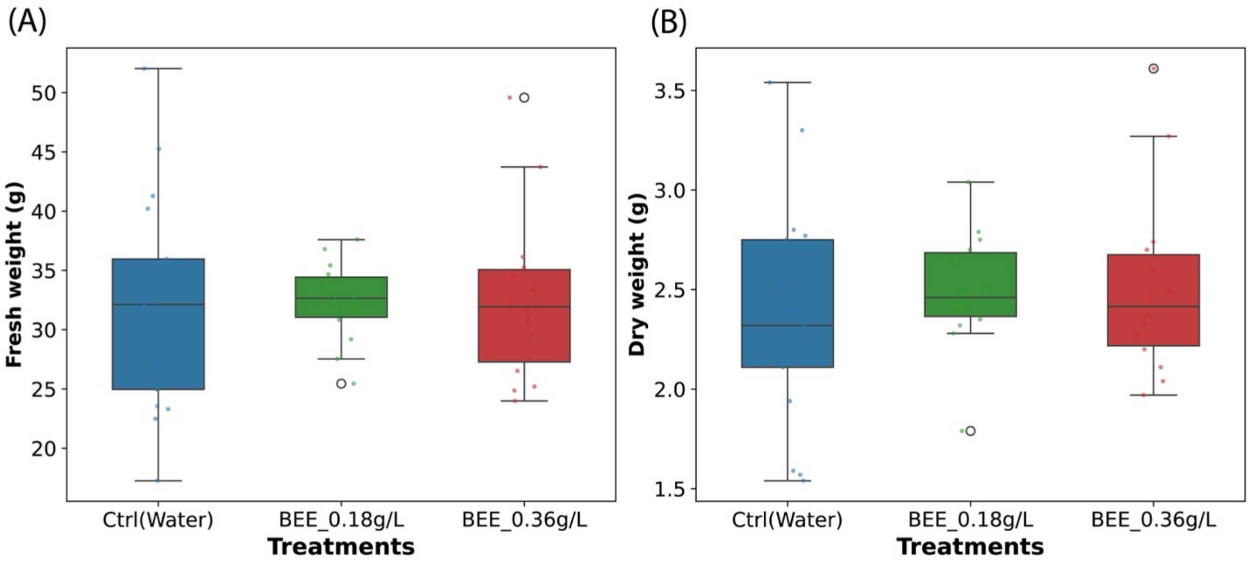


Figure S4. Graphical representation of the effect of BEE treatment or not (Ctrl (Water)) on Lettuce fresh (A) and dry (B) weight. Graph is presented in a boxplot with all datapoints overlaid. N= 15-18.

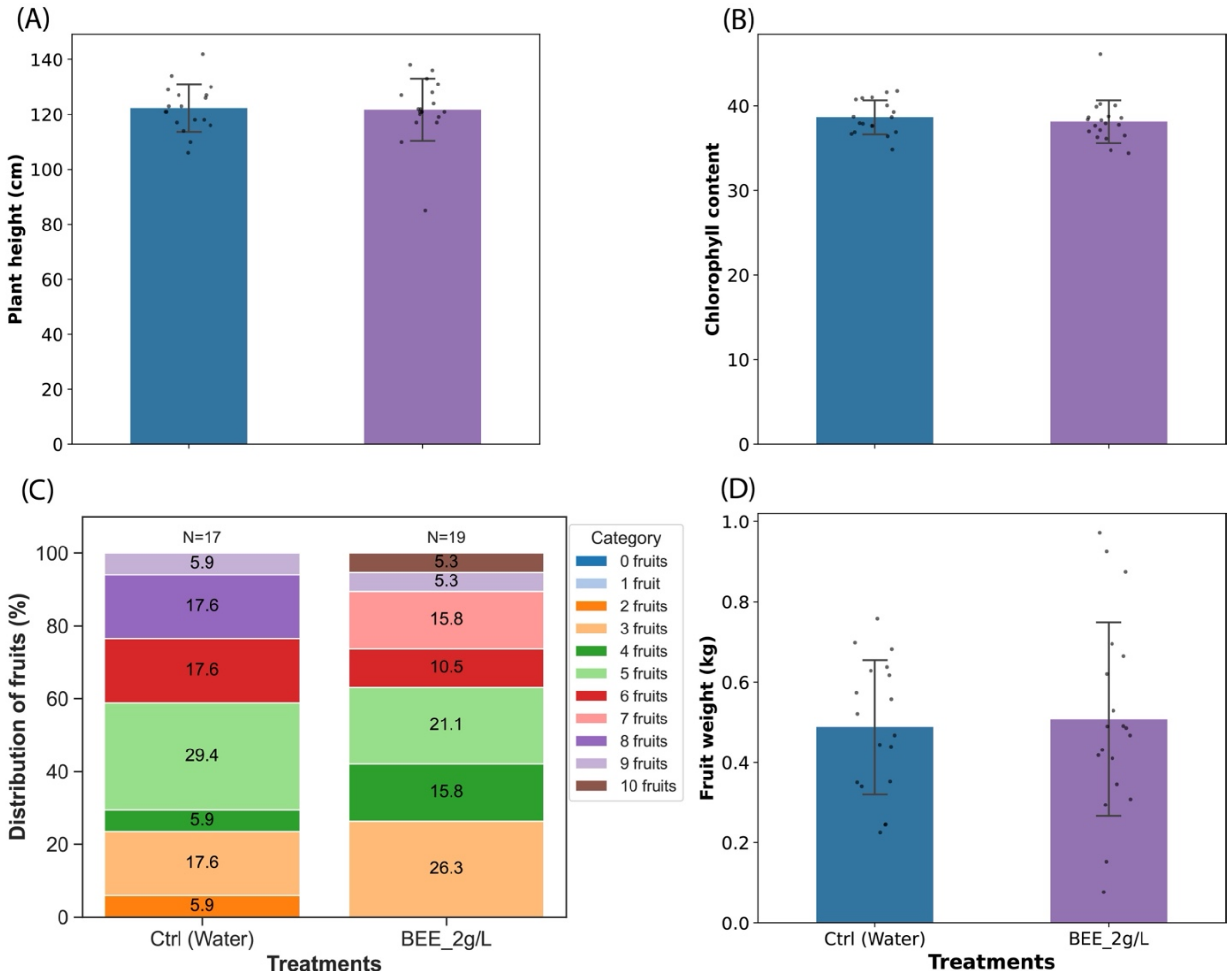


Figure S5. Effect of BEE treatment or not on growth parameters and yield of paprika plants under greenhouse conditions. (A) The average plant height (n=20-25). (B) The chlorophyll content from DUALEX readings (n=20-25). (C) The percentage distribution of fruits per treatment. (D) The total fresh weight of all fruits per treatment (n=20-25). Graphs are presented with error bars indicating standard deviation, and statistical significance (indicated by letters) was determined using ANOVA followed by Tukey's HSD test. The darker the colour of the bar in the plot, the more the mean. BEE: Belgian endive extract. Ctrl: Control treated with water.

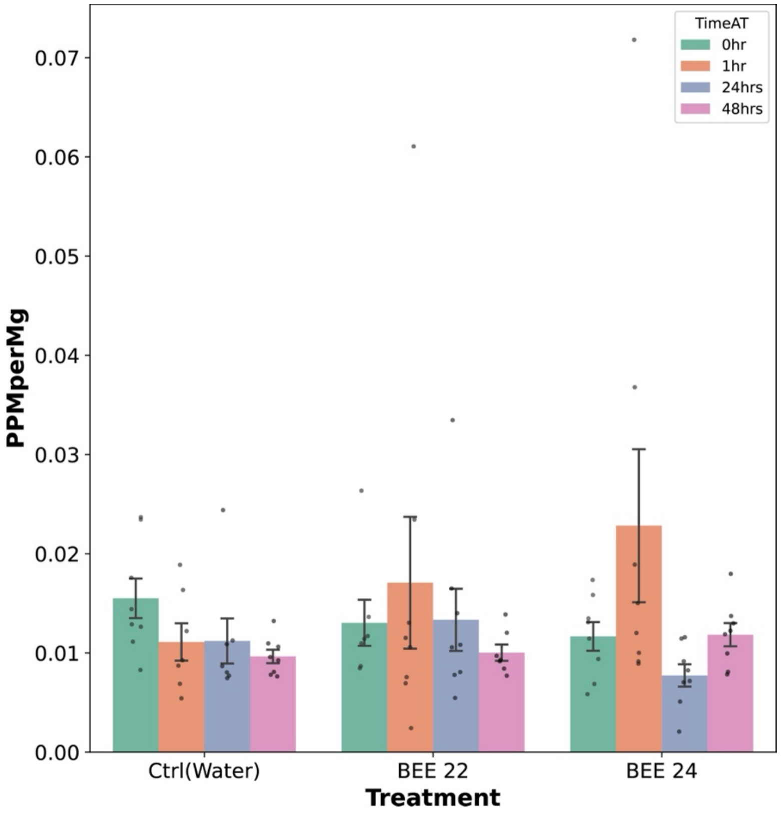


Figure S6. The ethylene quantified from plants treated with BEE or not (Ctrl) for 1 hour, 24 hours, and 48 hours. BEE: Belgian endive extract. Ctrl: Control treated with water. Graph is presented with error bars indicating standard deviation and statistical significance (indicated by p-values).

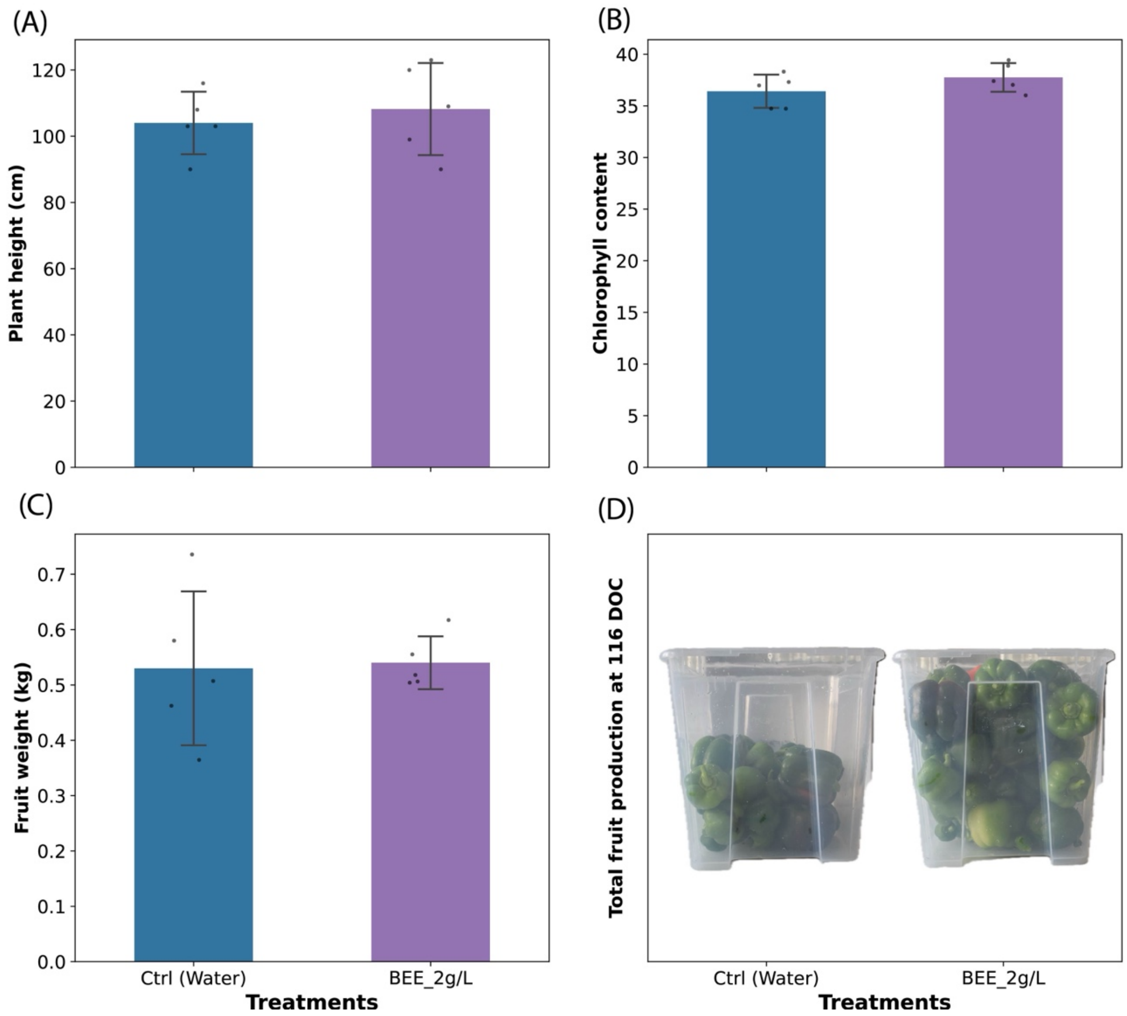


Figure S7. Effect of BEE treatment or not on growth parameters and yield of drought stressed sweet pepper plants under greenhouse conditions in 2022. (A) The average plant height. (n=5-10). (B) The chlorophyll content from DUALEX readings (n=5). (C) The total fresh weight of all fruits per treatment (n=5-10). (D) Pictures comparing the fruit size from treated and untreated groups. All graphs are presented with error bars indicating standard deviation and statistical significance was performed using t-test. BEE: Belgian endive extract. Ctrl: Control treated with water.

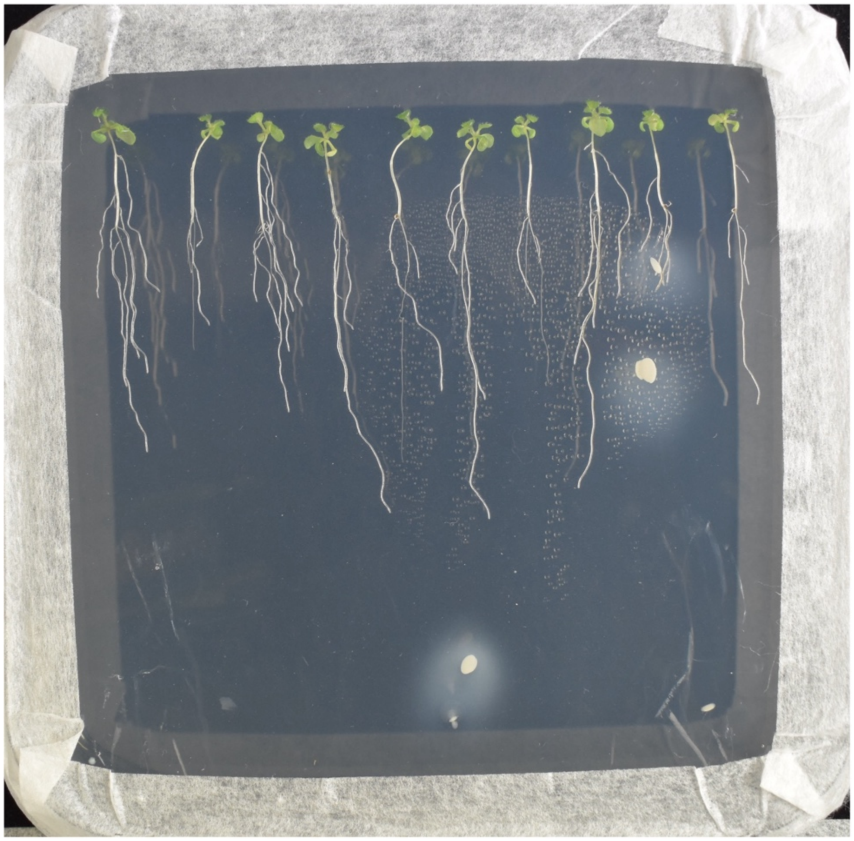

Figure S8. Picture of Arabidopsis seedlings treated with non-autoclaved BEE. The treatment caused contamination in the growth medium.

56
